# Secreted cytokines from inflammatory macrophages modulate sex differences in valvular interstitial cells on hydrogel biomaterials

**DOI:** 10.1101/2024.11.15.623805

**Authors:** Nicole E. Félix Vélez, Kristi Tu, Peng Guo, Ryan R. Reeves, Brian A. Aguado

**Affiliations:** Shu Chien-Gene Lay Department of Bioengineering, University of California, San Diego, La Jolla, CA, 92093, USA; Sanford Consortium for Regenerative Medicine, La Jolla, CA, 92037, USA; Nikon Imaging Center, Department of Cellular and Molecular Medicine, University of California, San Diego, La Jolla, CA, 92093, USA; Sulpizio Cardiovascular Center, University of California, San Diego, La Jolla, CA, 92093, USA; Program in Materials Science and Engineering, University of California, San Diego, La Jolla, CA, 92093, USA

**Keywords:** inflammation, sex differences, myofibroblasts, fibrosis, calcification, macrophages, hydrogels

## Abstract

Patients with aortic valve stenosis (AVS) experience fibrosis and/or calcification in valve tissue, which leads to heart failure if left untreated. Inflammation is a hallmark of AVS and secreted cytokines from pro-inflammatory macrophages are thought to contribute to valve fibro-calcification by driving the activation of valvular interstitial cells (VICs) to myofibroblasts. However, the molecular mechanisms by which inflammatory cytokines differentially regulate myofibroblast activation as a function of biological sex are not fully defined. Here, we developed an *in vitro* hydrogel culture platform to culture male and female valvular interstitial cells (VICs) and characterize the sex-specific effects of inflammatory cytokines on VIC activation to myofibroblasts and osteoblast-like cells. Our data reveal that tumor necrosis factor alpha (TNF-α) modulates female-specific myofibroblast activation via MAPK/ERK signaling, nuclear chromatin availability, and osteoblast-like differentiation via RUNX2 nuclear localization. Collectively, hydrogel biomaterials as cell culture platforms are critical for distinguishing sex differences in cellular phenotypes.

## Introduction

Sex differences in aortic valve stenosis (AVS) progression have been clinically documented, where male and female patients experience varied progression of aortic valve leaflet thickening and fibrocalcification.^1,2^ Specifically, male patients have higher valve calcification while female patients exhibit increased fibrotic remodeling, which can lead to sudden heart failure if left untreated.^3^ The current gold standard treatment for AVS is surgical or transcatheter aortic valve replacement; however, valve replacement patients are at risk of restenosis within a year of valve replacement surgery.^4^ Thus, small molecule drug treatments aimed at halting AVS or slowing its progression as an alternative for surgical intervention are needed. Moreover, since AVS progression is known to be heterogeneous between male and female patients,^5,6^ an opportunity exists to determine sex-specific treatments for AVS. Despite the clinical observations that AVS experience sex-specific disease progression,^3,5,7^ the underlying molecular mechanisms that drive sex dimorphisms in AVS pathogenesis continue to be poorly understood.

Valvular interstitial cells (VICs) are the resident fibroblasts in aortic valve tissue and regulate sex-specific tissue fibro-calcification via biochemical signaling and extracellular matrix (ECM) remodeling.^8^ As the valve tissue experiences routine hemodynamic stress, VICs transiently activate to myofibroblasts capable of remodeling their surrounding microenvironment and deactivate once homeostasis is achieved.^8,9^ During AVS progression, however, VICs remain persistently activated as myofibroblasts, resulting in aberrant ECM deposition, leading to pathological fibrosis.^9,10^ In AVS, VICs can also differentiate further to an osteoblast-like phenotype capable of depositing calcium phosphate, contributing to calcific nodule formation in the tissue.^8^ VICs are mechanosensitive cells, and matrix cues such as stiffness can regulate VIC activation into a myofibroblast phenotype.^11^ Sex differences in VICs have been previously observed in male and female VICs cultured on hydrogels mimicking the extracellular matrix properties of aortic valve leaflets. Aguado et al. identified increased myofibroblast activation in female VICs as a result of genes that escape X chromosome inactivation in stiff hydrogel cultures mimicking the diseased aortic valve tissue matrix.^12^ Given VICs have sex-specific mechanotransduction signaling pathways, we posit VICs have sex-specific responses to a wide array of microenvironmental cues.

Inflammation plays a critical role in valve fibro-calcification during AVS,^9,13–15^ and pro-inflammatory cytokines have been shown to drive sex differences in human VIC phenotypes, promoting an osteoblast-like phenotype in male VICs via sex-specific ERK pathway activation.^16^ Immune cells, such as pro-inflammatory macrophages, are known to infiltrate the aortic valve and secrete cytokines that help regulate VIC phenotype and matrix remodeling.^9,17^ One such cytokine, tumor necrosis factor alpha (TNF-α), has been previously implicated to both activate VICs and deactivate valve myofibroblasts, as well as exhibit varying calcifying effects in aortic valves.^18–20^ Moreover, high levels of TNF-α were observed in AVS patient sera after aortic valve replacement, with male patients exhibiting higher abundancy of TNF-α in their sera^20^, which may be linked to valve re-stenosis in male patients. Our understanding of the competing effects of TNF-α on fibro-calcification could be improved with a systematic analysis of sex-specific cytokine effects on VIC phenotype.

Here, we investigate the role of inflammatory cytokines, with a specific emphasis on TNF-α, in regulating sex differences in the transition states between quiescent VICs, activated myofibroblasts, and differentiated osteoblast-like cells. Using hydrogels as a cell culture platform recapitulating aortic valve matrix properties, we cultured male and female porcine VICs with conditioned media from M1 macrophages as well as physiologically relevant doses of TNF-α and evaluated the sex-specific changes in myofibroblast activation and osteoblast-like differentiation. We also investigated the role of MAPK/ERK signaling in mediating sex-specific TNF-α response, identifying a potential target for TNF-α-mediated AVS therapies. Collectively, our work suggests sex-specific regulation of MAPK/ERK signaling via TNF-α, which provides a bridge toward understanding sex-specific VIC phenotypes during AVS progression.

## Materials & Methods

### Conditioned Media Preparation and Characterization

M1 conditioned media was made as previously described.^9^ Briefly, THP-1 monocytes (ATCC, Cat. No. TIB-202) were thawed and cultured in RPMI media supplemented with 10% fetal bovine serum, 50 U/mL penicillin, 50 ug/mL streptomycin, and 1 ug/mL amphotericin B (THPI-1 media). Using a cell suspension of 1 million cells/mL, cells were centrifuged at 200 g for 10 minutes and resuspended in fresh THP-1 media supplemented with 100 ng/mL of phorbol myristate (PMA) (Sigma Aldrich, Cat. No. 79346) and incubated at 37°C with 5% oxygen in the dark for 24 hours. Cells were then centrifuged to remove media containing PMA and then incubated with fresh THP-1 media for 24 hours. Lipopolysaccharides (LPS, 10 ng/mL) and interferon-y (IFN-y, 50 ng/mL) were added and cultured for 48 hours to differentiate THP-1 cells to M1 macrophages. After differentiation, M1 macrophages were cultured with VIC media (Media 199 supplemented with 1% FBS, 1% penicillin/streptomycin, and 2% amphotericin B) for 48 hours. The M1 conditioned media was collected, centrifuged to remove cell debris, and stored at -80°C until use. The M1 conditioned media was characterized using Human Cytokine Array C5 (RayBiotech, Cat. No. AAH-CYT-5-2) following manufacturer protocols and normalized to VIC culture media. Chemiluminescent signals were analyzed using a Mini-Med x-ray film processor (AFP Manufacturing).

### Hydrogel Fabrication

Hydrogels were formed on 12 mm or 25 mm circular glass coverslips treated with (3-Mercaptopropyl)trimethoxysilane (MPTS, Sigma Aldrich, Cat. No. 175617) via overnight vapor deposition at 60°C. Poly(ethylene glycol) norbornene (PEG-Nb) was synthesized as previously described.^9,20,21^ Hydrogel precursor solution was made using a 4% w/v of 8-arm 40 kDa PEG-Nb mixed with 10% w/v 5 kDa PEG-dithiol (JenKem), 2 mM cysteine-arginine-glycine-aspartic acid-serine (CRGDS) cell adhesive peptide (Bachem) at a 0.99 thiol:ene ratio in PBS (Fisher Scientific, Cat. No. 14-190-250). Before polymerization, lithium phenyl-2,4,6-trimethylbenzoylphosphinate (LAP, Sigma-Aldrich, Cat. No. 900889-1G) was added at a 0.05 wt% in the dark. The gel precursor solution was pipetted onto a Sigmacote (Sigma-Aldrich, Cat. No. SL2-100ML) treated microscope slide and sandwiched by placing a thiolated glass coverslip on top. The hydrogel precursor solution was then photopolymerized using 365 nm light at 4 mW/cm^2^ for 3 minutes. The photopolymerized gels were sterilized in 5% isopropyl alcohol in PBS for 20 minutes, washed 3X with PBS, and swollen overnight in VIC culture media at 37°C and 5% CO2.

### Rheology

4% w/v PEG solution was photopolymerized *in situ* on a DHR 20 rheometer (TA Instruments). To characterize hydrogel formation, an 8 mm diameter geometry was used to perform oscillatory shear rheology with an amplitude of 1% and frequency of 1 Hz. The storage modulus obtained was converted to Young’s modulus (E) using the formular *E* = 2*G*^′^ · (1 + *v*), assuming a Poisson ratio of *v* = 0.5 since *G*^′^ >>> *G*′′.

### VIC Isolation and Expansion

VICs were isolated from fresh porcine hearts (Midwest Research) as previously described.^22^ Briefly, aortic valve leaflets were isolated from porcine hearts from 6-to-8 month-old adult pigs and rinsed with Earle’s Balanced Salt Solution (EBSS) (Sigma-Aldrich, Cat. No. E2888) supplemented with 5% penicillin/streptomycin, and 2% amphotericin B. Leaflets were then added to 250 units/mL type II collagenase (Fisher Scientific, Cat. No. NC9693955) solution in EBSS and incubated at 37°C under 5% CO2 for 30 minutes on a shaker at 100 rpm. Leaflets were then vortexed at maximum speed for 30 seconds and the remaining supernatant was removed. The leaflets were placed in fresh collagenase solution and incubated at 37°C under 5% CO2 for 60 minutes on a shaker at 100 rpm. The leaflets were then vortexed for 2 minutes at maximum speed to break them up. The remaining cell solution was passed through a 100 µm cell strainer. Cells were centrifuged at 200 g for 10 minutes and the cell pellet was resuspended in VIC expansion media (Media 199, 15% fetal bovine serum, 1% penicillin/streptomycin, and 2% amphotericin B). Cells were placed in T-75 cell culture flasks and incubated at 37°C under 5% CO2 for expansion. 70%-80% confluent VIC cultures were treated with trypsin for harvesting and counted using an automated cell counter (Countess).

### TNFα ELISA Characterization of M1 Conditioned Media and Human Serum

TNF-α enzyme linked immunoabsorbent assays (ELISA) were performed on M1 conditioned media and human serum (**Supplementary Table 1**) following the manufacturer’s protocol (Invitrogen, Cat. No. KHC3011). Serum samples were diluted ten-fold for the assay. Healthy samples were obtained from San Diego BloodBank. Diseased samples were obtained from AVS patients prior to transcatheter aortic valve replacement at Sulpizio Cardiovascular Center. All human serum was collected under approval from the UCSD Institutional Review Board (IRB #804209). All patients completed an IRB-approved consent form. All patient information has been de-identified for research purposes.

### Hydrogel VIC Culture

VICs were seeded on soft 2-dimensional PEG hydrogels at a density of 40,000 cells/cm^2^ in either VIC culture media (Media 199, 1% fetal bovine serum, 1% penicillin/streptomycin, and 2% amphotericin B) or M1 conditioned media for 3 days. Cells grown on 12- and 25-mm diameter gels were used for immunostaining experiments and quantitative reverse transcription (RT-qPCR), respectively. For TNF-α experiments, VICs were cultured on VIC culture media supplemented with 1 and 10 ng/mL of TNF-α and cultured for 3 days. For MAPK/ERK inhibition experiments, cells were treated with 10 µM of selumetinib (Selleck Chemical, Cat. No. AZD6244) solubilized in dimethyl sulfoxide (DMSO) at Day 0.

### Immunostaining and Immunofluorescent Characterization

After 3-day culture on hydrogels and TCPS, cells were fixed with 4% (v/v) paraformaldehyde in PBS (Electron Microscopy Sciences, Cat. No. 15710) for 20 minutes. Cells were then permeabilized with 0.1% (v/v) Triton-X 100 (Sigma Aldrich, Cat. No. 93443) in PBS for one hour. The samples were then blocked using 5% (w/v) bovine serum albumin (BSA, Sigma Aldrich, Cat. No. A8327) in PBS for one hour at room temperature. The samples were then stained with the following primary antibodies in blocking buffer and incubated for one hour at room temperature: αSMA (1:300, Abcam, Cat. No. ab7817), RUNX2 (1:300, Abcam, Cat. No. ab23981), methylated lysine (MeK, 1:350, Novus Biologicals, Cat. No. NB600824), and acetylated lysine (AcK, 1:350, Abcam, Cat. No. ab190479). Samples were washed with 0.05% (v/v) Tween-20 (Sigma Aldrich, Cat. No. P1379) in PBS. The samples were then stained with secondary antibodies Alexa Fluor 488 goat anti-mouse (1:200, Thermo Scientific, Cat. No. A11001) and Alexa Fluor 647 goat anti-rabbit (1:200, Thermo Scientific, Cat. No. A21245) in blocking buffer and incubated for one hour at room temperature in the dark. Nuclei were stained with DAPI (1:500, Roche, Cat. No.10236276001) and cell cytoplasm was stained with HCS Cell Mask Orange (1:5000, Fisher Scientific, Cat. No. H32713). Sample coverslips were rinsed once with 0.05% (v/v) Tween-20 in PBS and once with PBS before transferring to a glass-bottom well plate for imaging on a Nikon Eclipse Ti-2-E using a 20X objective. Image fluorescence was quantified by modifying a publicly sourced MATLAB code.^23^ Alpha smooth muscle actin intensity was normalized to CellMask cytoplasmic intensity. RUNX2 nuclear localization was quantified by normalizing RUNX2 nuclear intensity to cytoplasmic intensity. Nuclear methylated lysine ratios were calculated by normalizing MeK nuclear intensity to DAPI intensity. Nuclear acetyl lysine ratios were calculated by normalizing AcK nuclear intensity to DAPI intensity.

### Chromatin Condensation Parameter (CCP) Analysis

Nuclei were imaged as described above using DAPI staining. For CCP analysis, a gradient-based Sobel edge-detection algorithm was modified^24,25^ and employed to measure edge density on individual nuclei per image based on DAPI segmentation.

### RNA Isolation and RT-qPCR

RNA was isolated using a RNeasy Micro Kit (Qiagen, Cat. No. 74104) at the 3-day culture timepoint following the manufacturer’s protocols. RNA concentration was quantified using a NanoDrop 2000 spectrophotometer. Samples were normalized to 0.1 ng/uL concentration for cDNA synthesis using iScript Reverse Transcription Supermix for RT-qPCR (Bio-Rad, Cat. No. 1708841) following manufacturer’s protocol. To measure relative gene expression, iQ SYBR Green Supermix (Bio-Rad, Cat. No., 1708882) was mixed with corresponding primers (**Table 1**). Cq values were determined using a CFX384 iCycler. Gene expression of target genes was normalized to housekeeping gene *RPL30* expression.

**Table 1:**
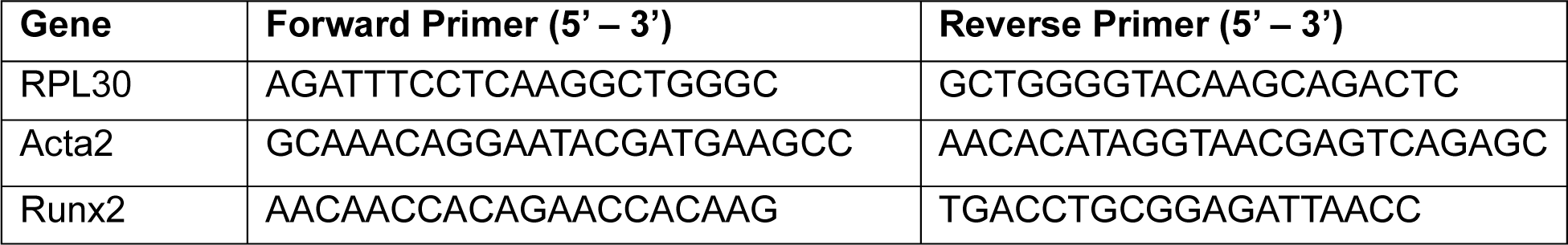
Primer sequences for RT-qPCR.

### Statistical Analysis

Statistical significance in immunofluorescence studies were determined using a two-way ANOVA with Tukey posttests and multiple comparisons in GraphPad PRISM. The threshold of significance was set to P<0.0001. Cohen’s d-test was performed between conditions and across sex to measure effect size for significant differences between groups that were determined to be statistically significant by multiple comparisons. Effect size significance was represented with the following key: *** = d > 0.8, ** = d > 0.5, * = d > 0.2 for between sex significance; ### = d > 0.8, ## = d > 0.5, # = d > 0.2 for male group significance; and $$$ = d > 0.8, $$ = d > 0.5, $ = 467 d > 0.2 for female group significance. For immunofluorescence studies, N=2 biological replicates and all data is presented as violin plots reporting median and first and third quartiles. For RT-qPCR studies, statistical significance was determined using a one-way ANOVA with Tukey posttests with significance threshold set to P<0.0001. The data for RT-qPCR studies is presented as individual technical replicates (n=5) across 2 biological replicates (N=2).

## Results

### Inflammatory factors from M1 macrophages drive VIC-to-osteoblast-like cell transition in female cells

Recognizing that AVS is a sex-specific, inflammation-mediated disease, we built upon prior work^9^ to characterize sex-specific VIC responses to secreted pro-inflammatory factors from M1 macrophages. We first sought to characterize our PEG hydrogel platform for cell culture. We leveraged thiol-ene click chemistry to form PEG hydrogels with cell-adhesive CRGDS peptides to recapitulate the stiffness of the healthy aortic valve matrix. Using shear rheology, we determined our hydrogels to have an elastic modulus of 5.38 ± 0.269 kPa, which were subsequently used for all hydrogel cultures (**Supplementary Figure 1**).

Using our hydrogels, we hypothesized that treatment with M1 pro-inflammatory factors would decrease myofibroblast activation in males relative to female VICs and increase osteoblast-like differentiation in males relative to females (**Fig. 1A**). Using αSMA stress fiber formation as a marker of myofibroblast activation, we observed female VICs exhibited increased activation than male VICs when cultured for 3 days with 1% FBS VIC culture media (control) in both hydrogel and tissue culture polystyrene (TCPS) cultures (**Fig. 1B,C**). We also found that after 3-day treatment with M1 conditioned media, both male and female VICs cultured on hydrogels exhibited an 80% reduction αSMA stress fiber formation relative to their respective male and female control groups, but female VICs had increased αSMA relative to males after M1 treatment, supporting our initial hypothesis (**Fig. 1C**). We also found that on TCPS, sex differences in VICs after M1 treatment were not significant, demonstrating hydrogels are critical for characterizing sex-specific cellular phenotypes (**Fig. 1C**). We also observed no changes in αSMA stress fibers in males after M1 treatment relative to female VICs cultured with VIC culture media on TCPS. We also note 2-fold higher activation levels for male and female VICs cultured on TCPS relative to hydrogel-VIC cultures. We also evaluated changes in *Acta2* gene expression which codes for αSMA in male and female VICs cultured on hydrogels as a function of M1 media treatment. We found that *Acta2* gene expression was reduced after M1 treatment in both male and female VICs (**Supplementary Fig. 2A**), further corroborating our immunocytochemistry observations.

**Figure 1:**
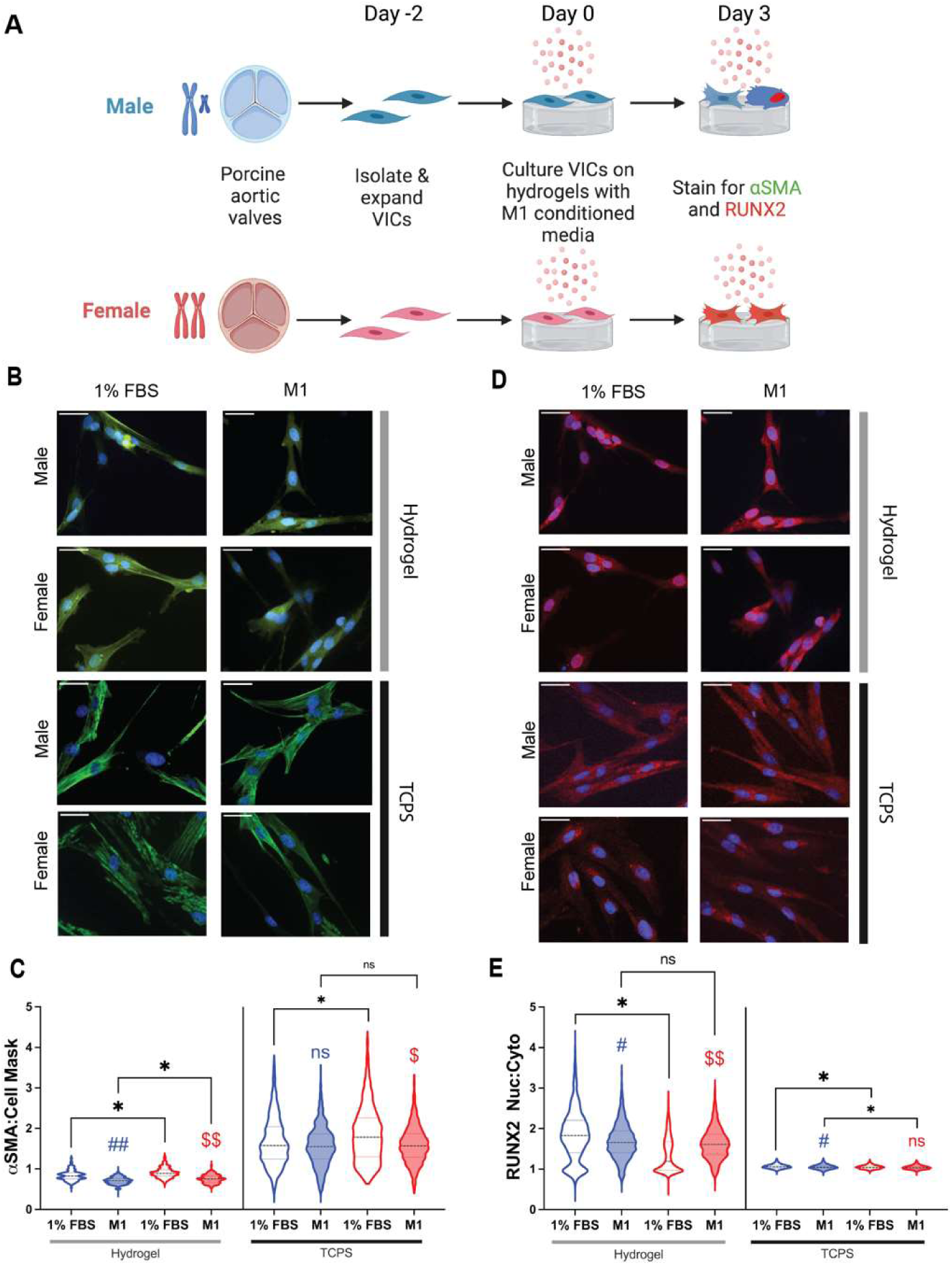
Treatment with M1 conditioned media decreases VIC myofibroblast activation and increases female-specific osteoblast-like differentiation. (**A**) Schematic of cell culture experiment to evaluate male and female porcine VIC phenotype via immunocytochemistry of alpha-smooth muscle actin (αSMA) and runt-related transcription factor 2 (RUNX2). (**B**) Representative images of male and female porcine VICs cultured on hydrogels or TCPS stained for alpha smooth muscle actin stress fibers (green) and nuclei (blue). Scale bar = 50 µm. (**C**) Quantification of αSMA intensity ratios in male (blue) and female (red) VICs treated with VIC culture media (1% FBS) and M1 conditioned media. (**D**) Representative images of male and female porcine VICs stained for RUNX2 (red) and nuclei (blue). Scale bar = 50 µm. (**E**) Quantification of RUNX2 nuclear localization in male (blue) and female (red) VICs treated with VIC culture media (1% FBS) and M1 conditioned media. For all graphs, data is reported for N=2 biological replicates and n=3 technical replicates. # and $ indicate statistical significance relative to same-sex control. * indicates sex difference. Statistical significance was determined by two-way ANOVA with Tukey posttests (P<0.0001) and effect size was measured between statistically significant groups using Cohen’s d test where *** = d > 0.8, ** = d > 0.5, * = d > 0.2.

Next, using RUNX2 nuclear localization as a marker of osteoblast-like differentiation, we observed male VICs treated with 1% FBS VIC culture media exhibited increased RUNX2 nuclear localization relative to female VICs in both hydrogel and TCPS cultures (**Fig. 1D,E**). In our hydrogel cultures, treatment with M1 conditioned media increased RUNX2 nuclear localization only in females, while surprisingly, males exhibited a slight decrease after M1 treatment relative to same-sex culture with VIC media (**Fig. 1E**). Gene expression of *Runx2* in hydrogel-VIC cultures after M1 treatment remained unchanged for male VICs relative to male control but increased in female VICs relative to female control (**Supplementary Fig. 2B**). Male VICs cultured on TCPS still exhibited a slight decrease in RUNX2 nuclear localization, but the effects of M1 on RUNX2 in female VICs was lost (**Fig. 1E**). Collectively, our results suggests that treatment of female VICs with M1 secreted factors increases RUNX2 nuclear localization only in female VICs cultured on hydrogels. Our data also suggests that our hydrogel cultures can maintain VIC quiescence relative to the TCPS cultures to better investigate the sex-specific effects of M1 pro-inflammatory factors on VIC phenotypes.

### Pro-inflammatory secreted factors modulate epigenetic changes in female VICs

We next hypothesized that M1 pro-inflammatory factors modulate female-specific changes in VIC chromatin structure to induce osteoblast-like differentiation. Previous work has demonstrated that myofibroblasts from diseased aortic valves exhibit decreased nuclear roundness relative to cells from healthy valves.^25^ We observed decreased nuclear roundness in both male and females VICs cultured on hydrogels treated with M1 when compared to cells cultured with VIC media (**Fig. 2A,B**). On the other hand, no changes in male nuclei roundness were observed in the TCPS cultures, but a decrease in nuclear roundness was observed in female M1-treated VICs grown on TCPS (**Fig. 2B**). We also note reductions in nuclear roundness were more pronounced in hydrogels relative to TCPS, with VICs on hydrogels exhibiting a 17% and 9.4% decrease in roundness for females and males respectively after M1 conditioned media treatment. VICs on TCPS only exhibited a 6.0% and 0.98% decrease in roundness after M1 conditioned media treatment (**Fig. 2B**).

**Figure 2:**
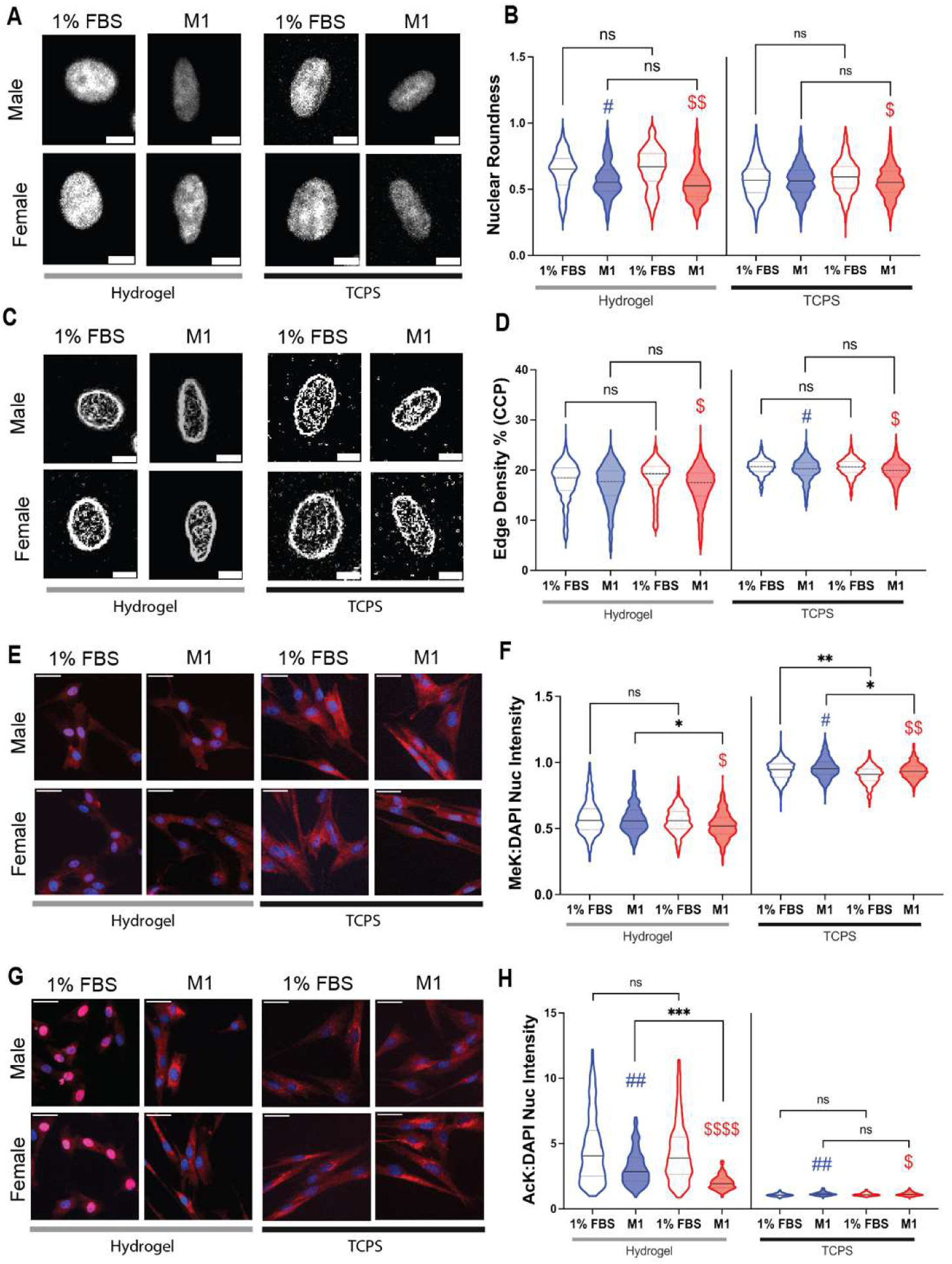
Effects of pro-inflammatory cytokines from M1 conditioned media on male and female VIC chromatin structures. (**A**) Greyscale images of single DAPI-stained nuclei. Scale bar = 10 µm. (**B**) Quantification of nuclear roundness in male (blue) and female (red) VICs treated with VIC culture media (1% FBS) and M1 conditioned media. (**C**) Representative images of single nuclei processed with a Sobel edge filter to calculate edge density percentage (CCP). (**D**) Quantification of edge density percentage to determine chromatin condensation. (**E**) Representative images of methylated lysine (red) and nuclei (blue). Scale bar = 50 µm. (**F**) Quantification of nuclear methylated lysine normalized to DAPI. (**G**) Representative images of acetylated lysine (green) and nuclei (blue). Scale bar = 50 µm. (**H**) Quantification of nuclear acetylated lysine normalized to DAPI. For all graphs, data is reported for N=2 biological replicates and n=3 technical replicates. # and $ indicate statistical significance relative to same-sex control. * indicates sex difference. Statistical significance was determined by two-way ANOVA with Tukey posttests (P<0.0001) and effect size was measured between statistically significant groups using Cohen’s d test where *** = d > 0.8, ** = d > 0.5, * = d > 0.2.

Next, we quantified chromatin condensation parameter (CCP) of DAPI-stained nuclei as an indicator of chromatin availability. When cultured on hydrogels, only female VICs treated with M1 media had decreased chromatin condensation relative to female VICs treated with control VIC media (**Fig. 2C,D**). Both male and female M1-treated VICs cultured on TCPS exhibited a decrease in chromatin condensation relative to same-sex groups treated with VIC media (**Fig. 2D**). Recognizing that histone acetylation promotes chromatin accessibility and histone methylation reduces chromatin accessibility, we characterized methylation and acetylation states in VICs treated with M1 conditioned media. We observed a female-specific decrease in methylation in our hydrogel-VIC cultures treated with M1 media relative to female cells treated with VIC media (**Fig. 2E,F**), consistent with female-specific decrease in CCP (**Fig. 2D**). On TCPS, we observed increased methylation in both male and female VICs relative to same-sex cells cultured with VIC media (**Fig. 2E,F**). Interestingly, we found decreased acetylation within the nucleus for male and female VICs cultured on hydrogels relative to cells treated with VIC media (**Fig. 2G,H**), while on TCPS, both male and female cells exhibited increased nuclear acetylation when treated with M1 media (**Fig. 2G,H**). Our results indicate that our hydrogel-VIC cultures treated with M1 pro-inflammatory factors recapitulate nuclear roundness associated with disease than TCPS. We also found that M1 proinflammatory factors promotes female-specific chromatin accessibility which may enable VIC-to-osteoblast-like cell differentiation.

### TNF-α drives female-specific osteoblast-like differentiation

We also identified candidate drivers of female-specific osteoblast-like differentiation within the M1 conditioned media. We performed a human cytokine array on the M1 conditioned media and the top three most abundant factors in M1 media include tumor necrosis factor alpha (TNF-α), interleukin 8 (IL-8), and regulated on activation, normal T cell expressed and secreted (RANTES) (**Fig. 3A, Supplementary Fig. 3A,B**) relative to the 1% FBS VIC culture media. Using an enzyme-linked immunosorbent assay (ELISA), we found that the M1 media has a TNF-α concentration of 0.125 ng/mL (**Supplementary Fig. 4**). We compared TNF-α levels in M1 conditioned media to levels found on human AVS patient sera using an ELISA. We found that male and female AVS patient serum contained 90.05 ± 12.98 pg/mL and 87.87 ± 15.75 pg/mL of TNF-α, respectively (**Fig. 3B**).

**Figure 3:**
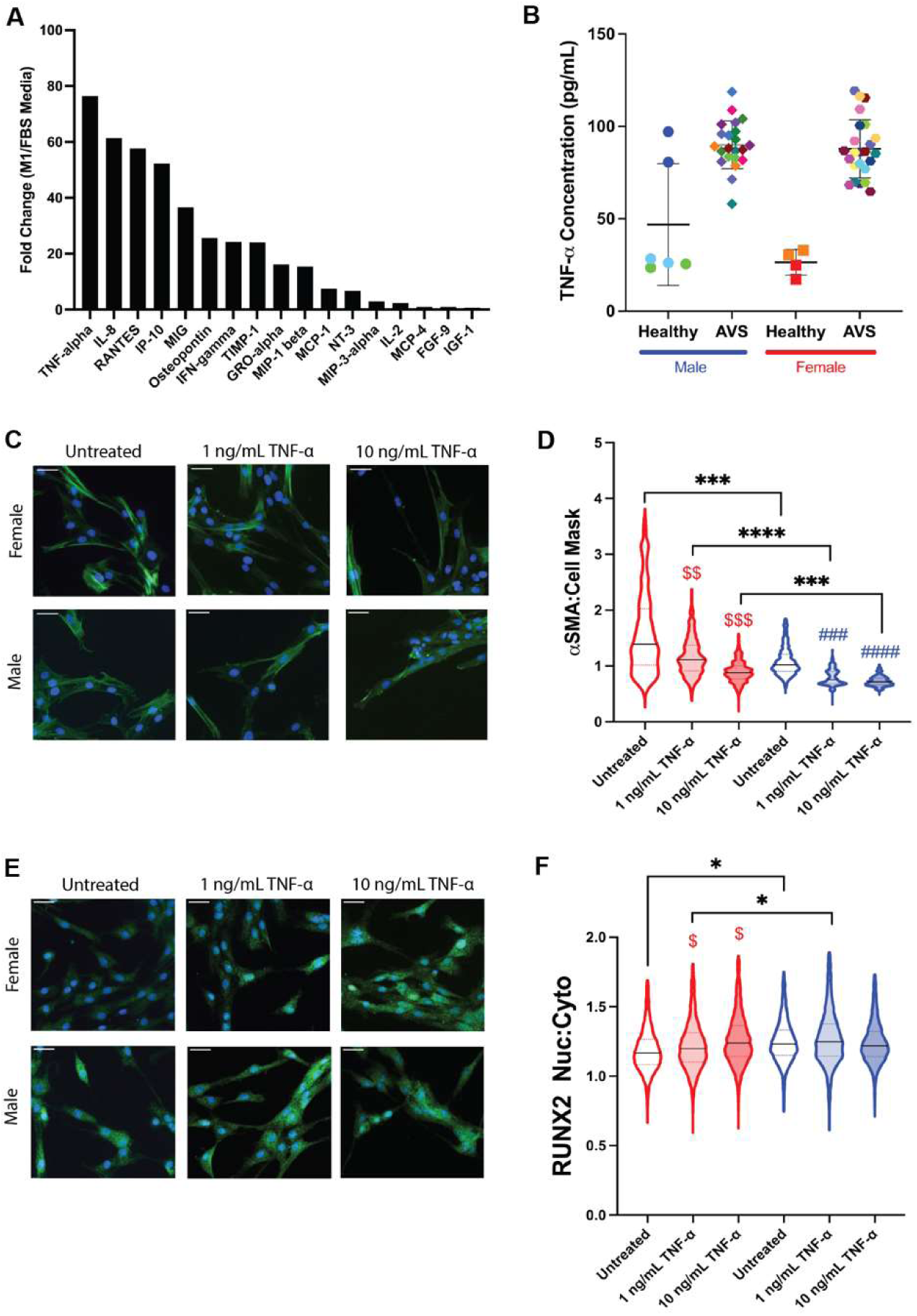
TNF-α drives sex-specific osteoblast-like differentiation in female VICs. (**A**) Human cytokine array on M1 conditioned media relative to VIC culture media. (**B**) Quantification of TNF-α in human serum. N=4 healthy male and female patients and N=22 male AVS patients and N=24 female AVS patients. Colors indicate individual patients. (**C**) Representative images of male and female porcine VICs cultured on hydrogels or TCPS stained for alpha smooth muscle actin stress fibers (green) and nuclei (blue). Scale bar = 50 µm. (**D**) Quantification of αSMA intensity ratios in female (red) and male (blue) VICs treated with VIC culture media (untreated), 1 ng/mL TNF-α, and 10 ng/mL TNF-α. (**E**) Representative images of male and female porcine VICs stained for RUNX2 (green) and nuclei (blue). Scale bar = 50 µm. (**F**) Quantification of RUNX2 nuclear localization in female (red) and male (blue) VICs treated with VIC culture media (untreated), 1 ng/mL TNF-α, and 10 ng/mL TNF-α. For all graphs, data is reported for N=2 biological replicates and n=3 technical replicates. # and $ indicate statistical significance relative to same-sex control. * indicates sex difference. Statistical significance was determined by two-way ANOVA with Tukey posttests (P<0.0001) and effect size was measured between statistically significant groups using Cohen’s d test where *** = d > 0.8, ** = d > 0.5, * = d > 0.2.

To further our understanding of how TNF-α mediates VIC phenotypes, we cultured male and female VICs on hydrogels with higher doses of 1 ng/mL and 10 ng/mL of TNF-α, recognizing TNF-α has a half-life of 15-30 minutes in serum.^26^ We observed that TNF-α decreased αSMA stress fiber formation in both males and females, but female VICs still maintain higher activation levels relative to males even after TNF-α treatment (**Fig. 3B,C**). TNF-α treatment increased RUNX2 nuclear localization uniquely in females relative to the female untreated group, while no changes were observed in males VICs treated with TNF-α relative to same-sex untreated group (**Fig. 3D,E**). Taken together, this data suggests that TNF-α uniquely drives female-specific osteoblast-like differentiation *in vitro*.

### Female VICs are more responsive to TNF-α signaling than male VICs

Since we observed similar female-specific changes in VIC phenotype as the M1 treated cultures, we hypothesized TNF-α induced female-specific chromatin relaxation. We quantified CCP after TNF-α treatment, and interestingly, we found that TNF-α itself does not cause changes to the chromatin structure relative to same sex untreated groups (**Fig. 4A,B**). We do note that TNF-α treatment did induce sex differences in CCP, with TNF-α-treated female VICs exhibiting decreased chromatin condensation relative to TNF-α-treated males (**Fig. 4A,B**). Next, we quantified nuclear methylation and found that nuclear methylation significantly decreased in both female and male VICs treated with TNF-α relative to same-sex untreated groups, with females exhibiting an increased fold change in methylation relative to same-sex control (**Fig. 4C,D**). We also quantified nuclear acetylation after TNF-α treatment and observed significantly increased nuclear acetylation in female VICs after treatment with both 1 ng/mL and 10 ng/mL of TNF-α relative to female untreated groups, while male VICs expressed increased nuclear acetylation only after treatment with 10 ng/mL TNF-α (**Fig. 4E,F**). Overall, these results suggests that female VICs are more responsive to TNF-α than males, and that TNF-α alone can cause female-specific chromatin relaxation relative to males treated with TNF-α.

**Figure 4:**
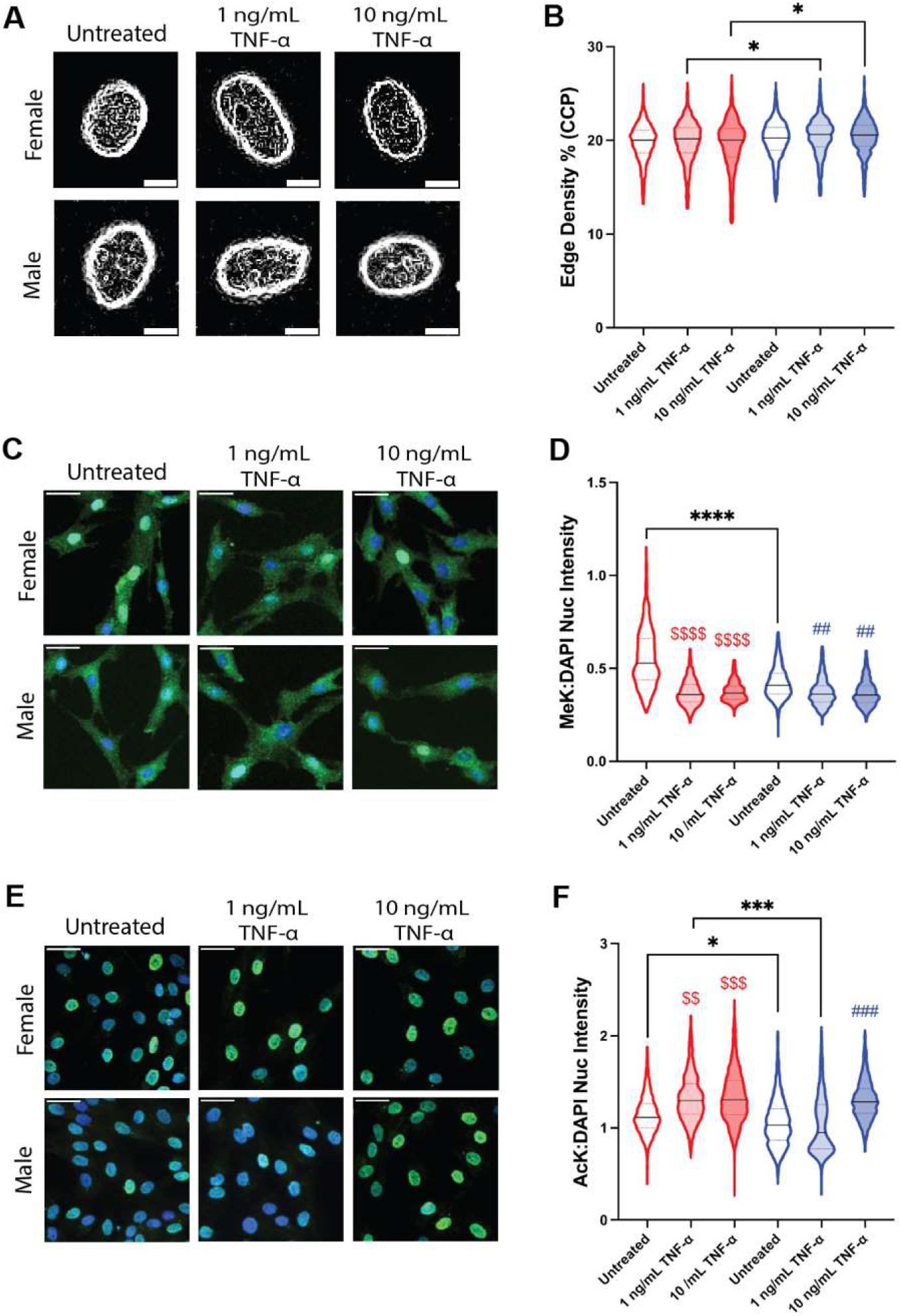
Effects of TNF-α from M1 conditioned media on male and female VIC chromatin structures. (**A**) Representative images of single nuclei processed with a Sobel edge filter to calculate edge density percentage (CCP). (**B**) Quantification of edge density percentage to determine chromatin condensation. (**C**) Representative images of methylated lysine (green) and nuclei (blue). Scale bar = 50 µm. (**F**) Quantification of nuclear methylated lysine normalized to DAPI. (**G**) Representative images of acetylated lysine (green) and nuclei (blue). Scale bar = 50 µm. (**H**) Quantification of nuclear acetylated lysine normalized to DAPI. For all graphs, data is reported for N=2 biological replicates and n=3 technical replicates. # and $ indicate statistical significance relative to same-sex control. * indicates sex difference. Statistical significance was determined by two-way ANOVA with Tukey posttests (P<0.0001) and effect size was measured between statistically significant groups using Cohen’s d test where *** = d > 0.8, ** = d > 0.5, * = d > 0.2.

### Inhibition of MEK 1/2 reverses TNF-α-mediated changes to female VIC phenotypes

Since the MAPK/ERK pathway is known to mediate TNF-α signaling,^19^ we hypothesized that inhibiting ERK signaling using the MEK 1/2 inhibitor selumetinib would prevent TNF-α-mediated osteoblast-like differentiation in female VICs. Using a TNF-α dose of 1 ng/mL, we found that selumetinib prevented TNF-α-mediated deactivation uniquely in female VICs relative to TNF-α treatment (**Fig. 5A**). We observed no difference between the TNF-α and TNF-α + selumetinib treatment groups in male VICs (**Fig. 5A**). In most treatment groups, female VICs still exhibited higher levels of αSMA stress fibers than males (**Fig. 5B**). We also observed that treatment with selumetinib reversed TNF-α-mediated osteoblast-like differentiation in female VICs relative to TNF-α (**Fig. 5C**). As before, male VICs demonstrated no significant difference between the TNF-α and TNF-α + selumetinib treatment groups (**Fig. 5C**). We found that male VICs had higher levels RUNX2 nuclear localization for untreated, vehicle control, and TNF-α + selumetinib groups, but treatment with TNF-α only in female VICs increased RUNX2 nuclear localization to ratios similar to the male TNF-α treated group (**Fig. 5D**). Overall, we observed that female VICs uniquely responded to MEK 1/2 inhibition, with MEK 1/2 inhibition preventing any changes in VIC phenotype after TNF-α treatment.

**Figure 5:**
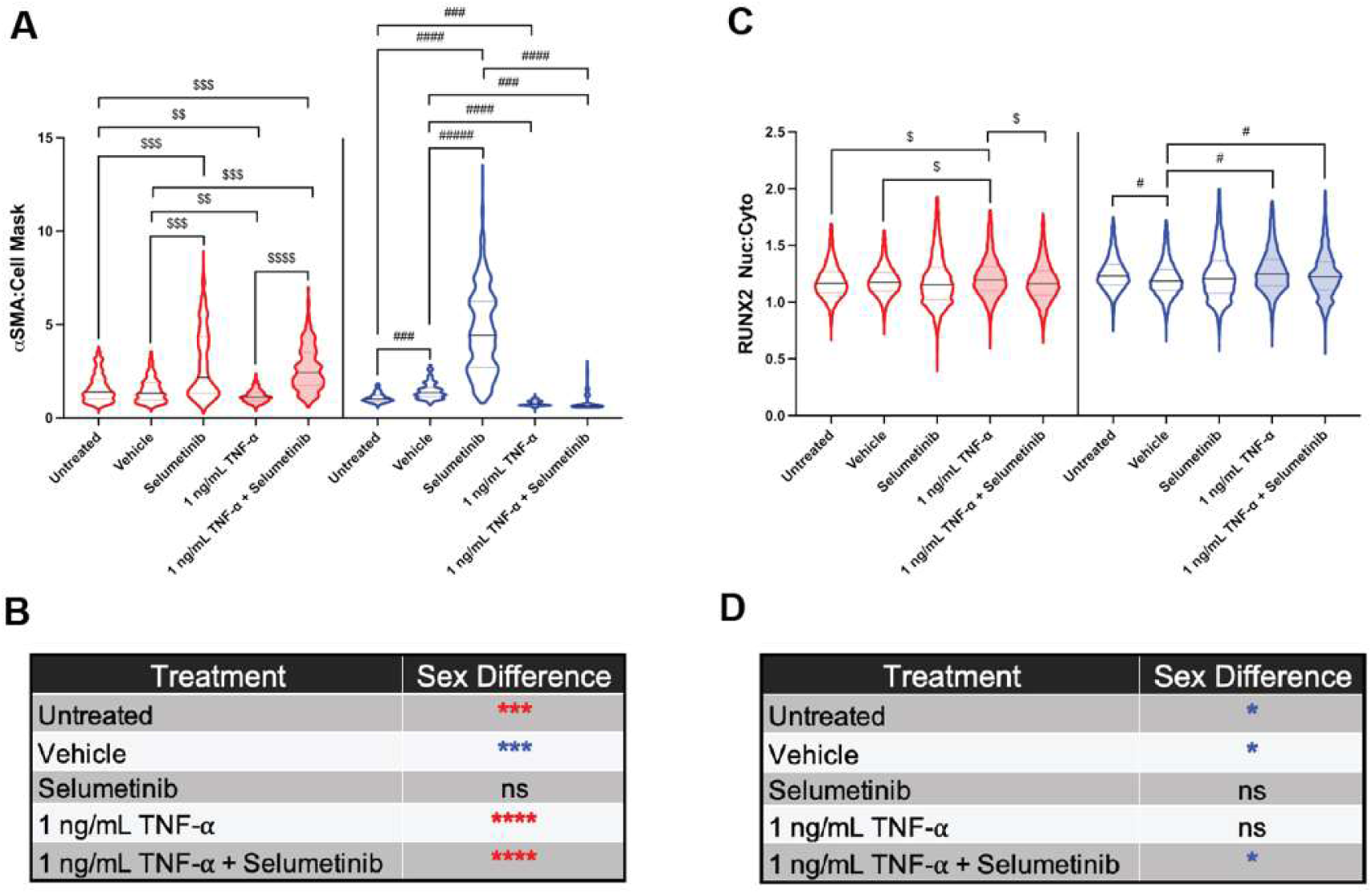
Inhibition of MAPK/ERK signaling with selumetinib reverses TNF-α-mediated changes to female VIC phenotypes. (**A**) Quantification of αSMA stress fibers relative to HCS Cell Mask in female (red) and male (blue) VICs cultured on hydrogels. (**B**) Tabulated chart showing sex differences between conditions in αSMA stress fiber quantification for each condition. (**C**) Quantification of RUNX2 nuclear localization in female (red) and male (blue) VICs cultured on hydrogels. (**D**) Tabulated chart showing sex differences between conditions in RUNX2 nuclear translocation for each condition. For all graphs, data is reported for N=2 biological replicates and n=3 technical replicates. Statistical significance was determined by two-way ANOVA with Tukey posttests (P<0.0001) and effect size was measured between statistically significant groups using Cohen’s d test where *** = d > 0.8, ** = d > 0.5, * = d > 0.2.

## Discussion

Our work here demonstrates that the pro-inflammatory factor TNF-α modulates female-specific VIC-to-osteoblast-like cell differentiation via MAPK/ERK signaling (**Fig. 6**). TNF-α is highly expressed in serum from patients with severe AVS relative to healthy samples^20,27^ (**Fig. 3B**), and previous work has demonstrated increased presence of M1 macrophages that secrete TNF-α in calcified aortic valve leaflets.^13^ The role of TNF-α in valvular fibro-calcification has long been debated, with some reports showing its antifibrotic effect^28^ while other demonstrated its calcifying effect.^18,29^ In these *in vitro* studies, myofibroblast cultures were performed in tissue culture polystyrene, a material with supraphysiological stiffness that automatically causes myofibroblast activation. Using hydrogels to promote fibroblast quiescence, previous work has shown in VICs that TNF-α has an antifibrotic effect on quiescent fibroblasts but plays a pro-calcific role in activated myofibroblasts.^19^ Building on this work, our contribution reveals female VICs have sex-dependent myofibroblast phenotypes in response to TNF-α. Specifically, we posit that female VICs interpret TNF-α and likely other pro-inflammatory cues via the MAPK/ERK signaling pathway to drive fibrosis and eventual calcification in the female aortic valve.

**Figure 6:**
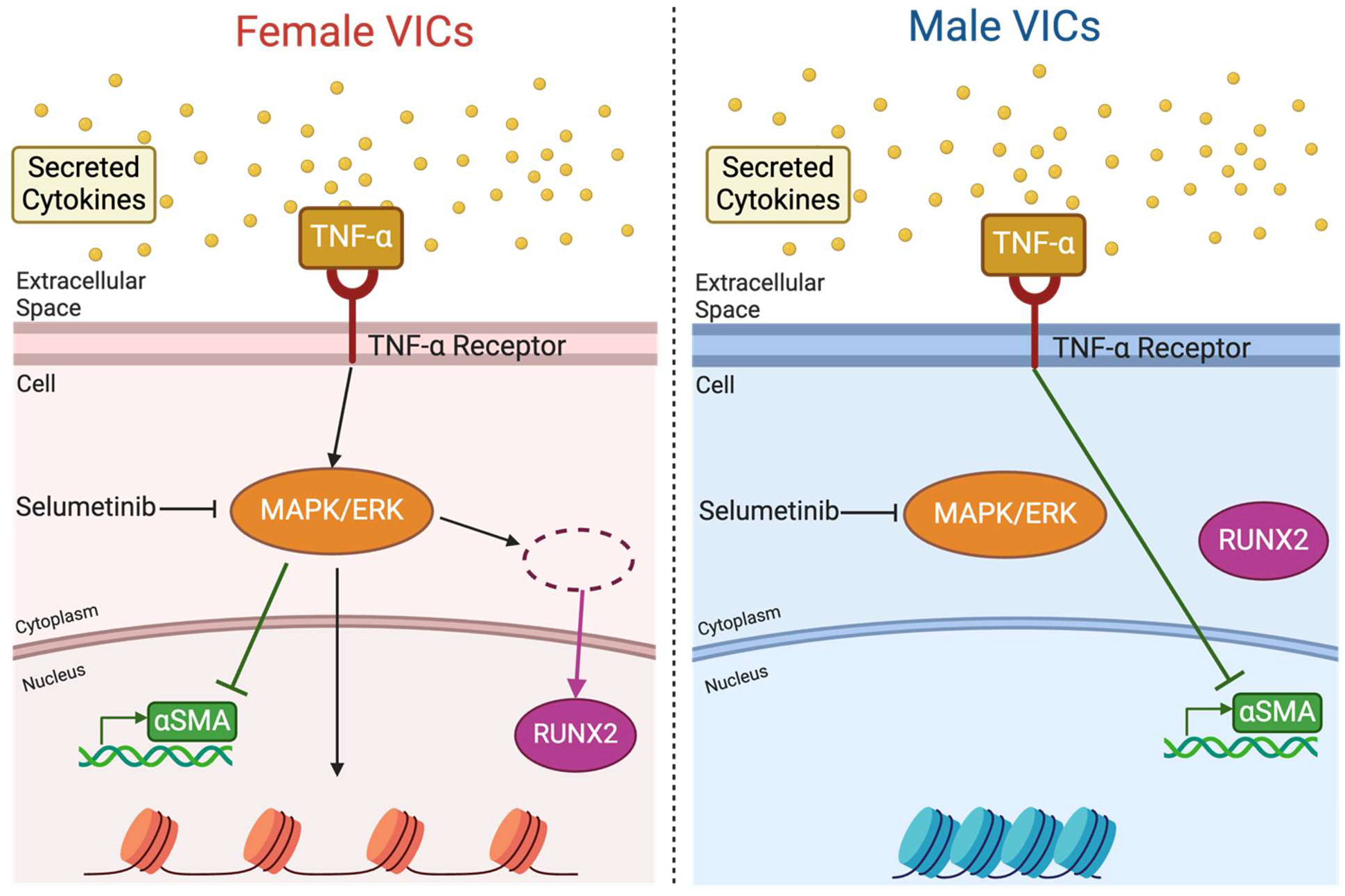
Proposed mechanism of action of TNF-α-mediated VIC-to-osteoblast-like cell differentiation. Female VICs exhibit decreased αSMA stress fiber formation and *Acta2* gene expression, increased RUNX2 nuclear localization, and more relaxed chromatin structure as a result of TNF-α signaling via MAPK/ERK pathway. Male VICs, on the other hand, only exhibited decreased αSMA stress fiber formation and *Acta2* gene expression as a response to TNF-α signaling.

We also reveal VICs undergo sex-specific epigenetic alterations in response to secreted inflammatory cytokines. By treating VICs with M1 secreted cytokines and physiologically relevant doses of TNF-α,^20,27^ we demonstrate that pro-inflammatory factors can modulate quiescent VIC phenotype in female cells by altering chromatin structures. We suggest that pro-inflammatory signaling can lead to female-specific chromatin relaxation that could then enable osteoblast-like cell differentiation. Epigenetic changes are a hallmark of aging, cardiovascular disease, and heart failure.^30,31^ In particular, chromatin remodelers such as histone acetyltransferases (HATs), histone deacetylases (HDACs), and histone demethylases act together to alter chromatin accessibility and consequently affect gene expression.^30^ Other work has demonstrated increased HAT activity in quiescent VICs relative to mechanically activated VICs, suggesting that during AVS, decreased HAT expression can contribute to AVS progression.^32^ Our work sets the stage for future studies to investigate how pro-inflammatory signaling leads to changes in chromatin remodeler activity and subsequent female VIC-to-osteoblast-like cell differentiation. Due to the female-specific results reported here, we posit X-linked histone demethylases such as lysine demethylase 5C (KDM5C) and lysine demethylase 6A (KDM6A), which are known to escape X chromosome inactivation, may modulate female-specific demethylation in response to pro-inflammatory cytokines.

Our work provides further evidence that hydrogel biomaterials as cell culture platforms are important tools to dissect sex-specific cellular phenotypes that recapitulate cell phenotypes in diseased aortic valve tissue. We demonstrated that our hydrogel cultures can maintain VICs in a quiescent cell state which helps to decouple the effects of biochemical signaling from mechanical cues. Similar work has shown that in valve and cardiac fibroblasts, biochemical signaling and stiffness cues have differing effects on myofibroblast protein expression,^20,23,32,33^ demonstrating that mechanical and biochemical cues have independent and synergistic effects on myofibroblasts. Previous work has also shown that nuclei in diseased aortic valve tissue are less round than nuclei in healthy tissue.^25^ In our work, we also demonstrate that treating cells cultured on hydrogels with pro-inflammatory factors can recapitulate important characteristics of disease such as decreased nuclei roundness in cells from diseased aortic valves. Leveraging our *in vitro* platforms, we suggest inflammatory factors during fibrosis modulates nuclear roundness and subsequent chromatin remodeling that occurs within the nuclei of VICs during AVS progression. Future studies that fully characterize sex-specific open chromatin regions and transcription factor binding sites available in response to inflammatory factors are warranted.

Hydrogel culture platforms will also become increasingly important to determine sex-specific cellular interactions with small molecule inhibitors. Using selumetinib, an MEK 1/2 small molecule inhibitor, we demonstrated a reversal of TNF-α-mediated calcification in females. Selumetinib is mostly used as a treatment for different types of cancers,^19,34–36^ but a preclinical study on a heart failure rat model revealed its potential as a therapeutic intervention for cardiac hypertrophy. Treatment with selumetinib was shown to prevent cardiac hypertrophy and fibrosis by preserving cardiomyocyte size and function after injury.^37^ Thus, further investigation into selumetinib as a therapeutic for aortic valve calcification is warranted. In order to accelerate sex-specific drug treatments for aortic valve disease, physiologically relevant *in vitro* cultures will help identify sex-specific targets for therapeutic intervention,^12^ as well as optimizing drug dosing using computational tools.^38–40^ We anticipate selumetinib will be an attractive small molecule inhibitor to target sex-specific AVS progression in future work.

There are several limitations to our current work. Firstly, our studies focused on paracrine signaling from M1 macrophages and did not consider intercellular interactions between VICs and macrophages. Future co-cultures with VICs and macrophages will help further understand how cell-cell interactions via cadherins can regulate sex-specific paracrine signaling effects and subsequent myofibroblast activation.^41^ Secondly, we focus on the role of TNF-α in female-specific VIC-to-osteoblast-like cell differentiation, while *in vivo* there are many other inflammatory factors acting in concert during the inflammatory stage of AVS.^9,20,42^ Moreover, our studies evaluated the effects of pro-inflammatory factors after three days in culture. Recognizing that AVS is a time-dependent disease, future studies should evaluate the spatial and temporal effects of pro-inflammatory signaling to determine more permanent phenotypical and epigenetic changes that arise from longer culture times with inflammatory factors and three-dimensional cell cultures.^43,44^

## Conclusions

In conclusion, our study suggests that pro-inflammatory factors and TNF-α can cause osteoblast-like cell differentiation in female VICs by inducing changes to the overall chromatin structure in the nucleus, which may contribute to female-specific fibro-calcification during AVS. We also highlight that TNF-α acts via MAPK/ERK signaling to induce female-specific phenotypic shifts. Using physiologically relevant *in vitro* cell culture platforms, we demonstrated that hydrogels mimicking aortic valve stiffness allow for controlled studies on VIC phenotype, separating biochemical and biomechanical influences. Our work also lays the foundation for future investigations into sex-specific inflammatory mechanisms in AVS progression. Moreover, our data point to selumetinib as a promising therapeutic candidate to reverse TNF-α-mediated calcification, warranting further exploration in the context of aortic valve disease.

## Acknowledgements

N.E.F.V. acknowledges funding from the National Science Foundation Graduate Research Fellowship Program (NSF-GRFP) and the UC San Diego GEMINI Graduate Fellowship. B.A.A. acknowledges support from the National Institutes of Health (R00 HL148542), the NIH Director’s New Innovator Award (DP2 HL173948), the Chan Zuckerberg Initiative Science Diversity Leadership Award, and the American Heart Association (942253). We thank all patients that provided serum samples for research purposes.

## Author Contributions

N.E.F.V. and B.A.A. conceived and supervised the study. N.E.F.V. and K.T. performed all *in vitro* experiments, analyzed data, and performed statistical analysis. P.G assisted in developing the MATLAB image analysis code. R.R.R. collected blood samples. N.E.F.V. and B.A.A. wrote and edited the manuscript. All authors approved the manuscript.

## Competing Interests

No disclosures to report.

## Supplementary Material

**Supplementary Figure 1:**
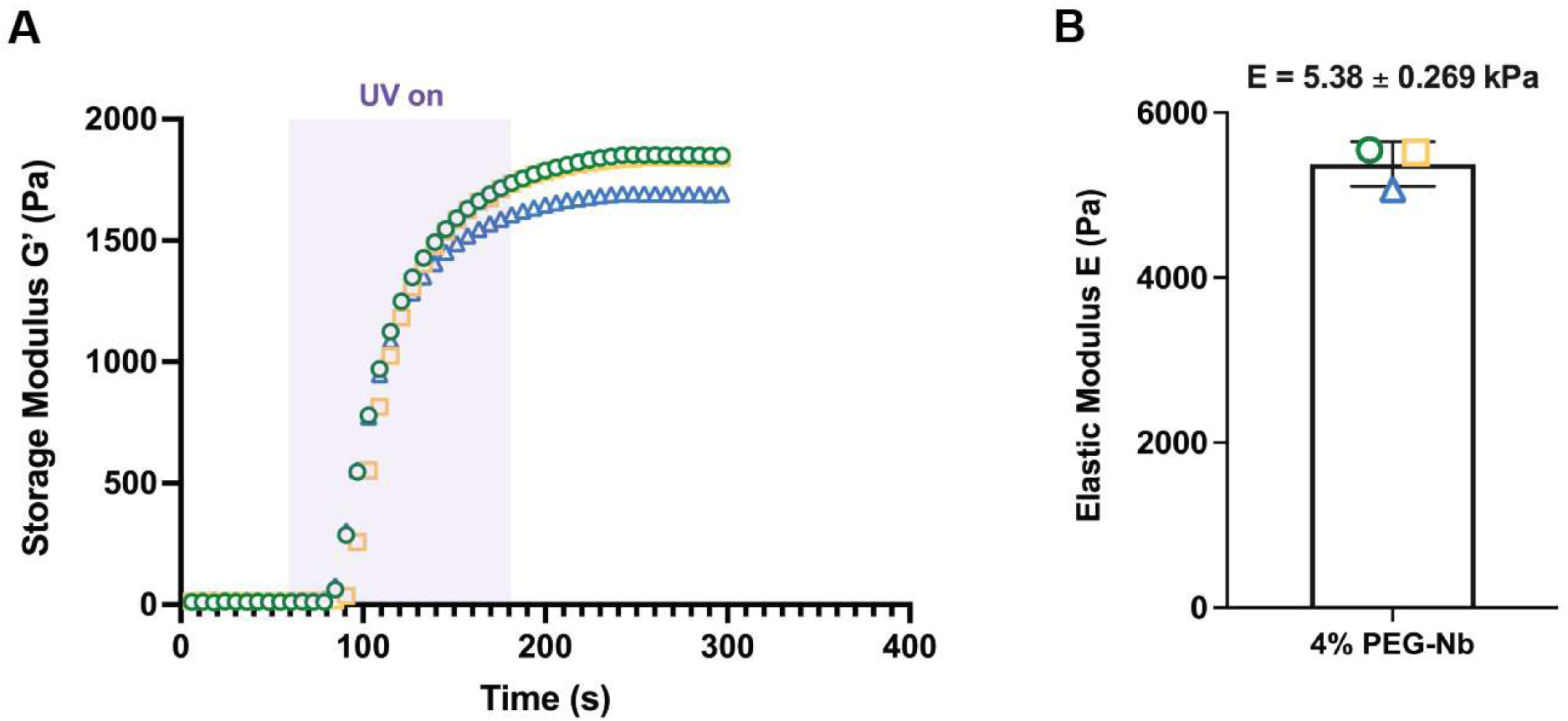
Rheological characterization of 4% PEG-Nb hydrogels. (**A**) Storage modulus of hydrogel solution upon exposure to UV light. (**B**) Final elastic modulus of hydrogel after UV light exposure with mean ± standard deviation shown.

**Supplementary Figure 2:**
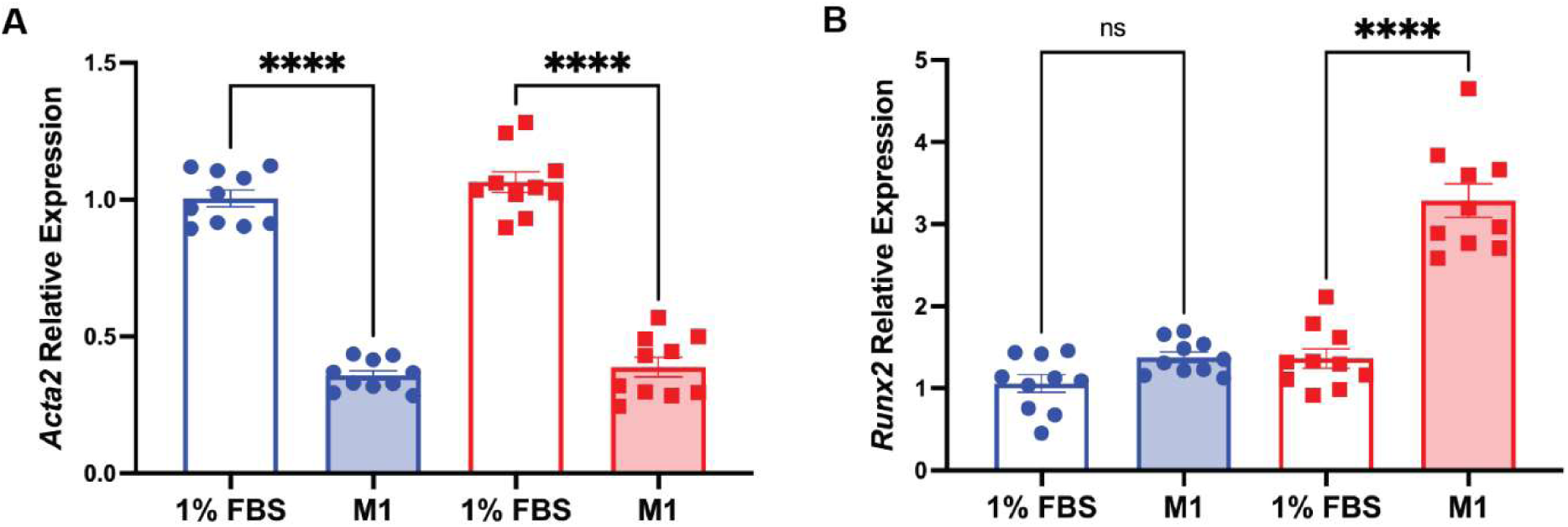
Quantitative *Acta2* and *Runx2* gene expression. (**A**) *Acta2* and (**B**) *Runx2* gene expression in male (blue) and female (red) VICs cultured on hydrogels treated with VIC culture media (1% FBS) and M1 conditioned media. Statistical significance was determined using a one-way ANOVA with Tukey posttests and indicated by ****=P<0.0001.

**Supplementary Figure 3:**
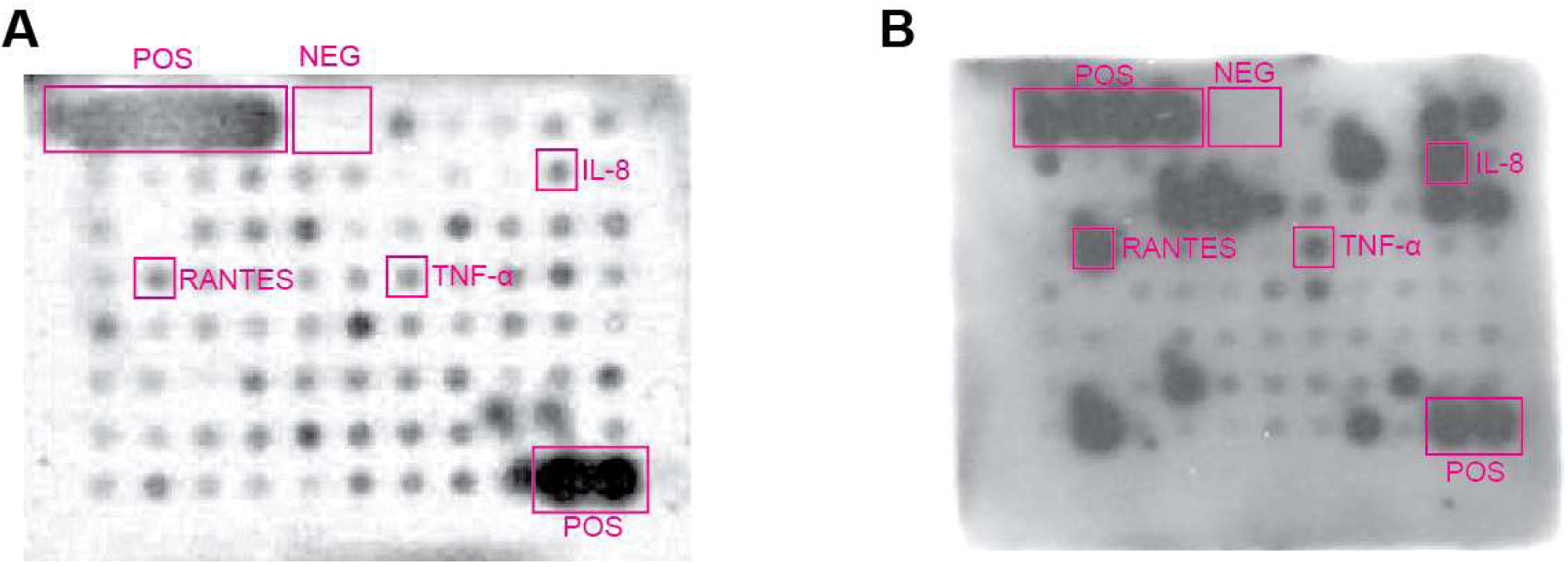
Cytokine array blots for (A) 1% FBS VIC culture media and (B) M1 conditioned media. The top 3 proteins (TNF-α, IL-8, and RANTES) with greatest fold change have been labeled with boxes (POS = manufacturer positive control, NEG = manufacturer negative control).

**Supplementary Figure 4:**
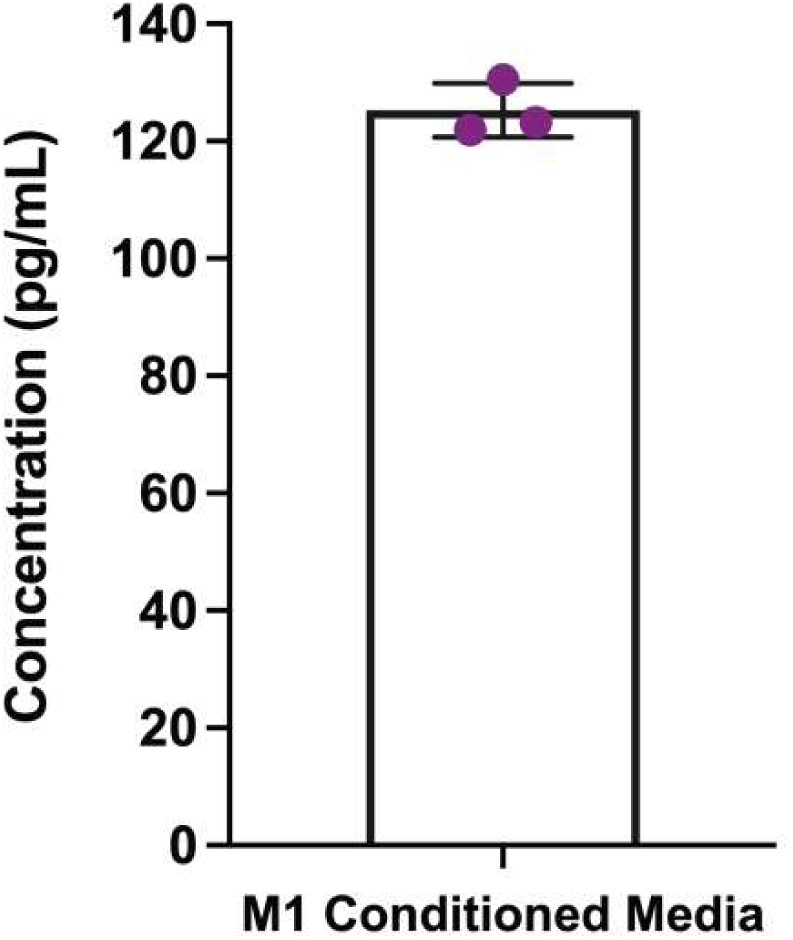
TNF-α ELISA of M1 conditioned media.

**Supplementary Table 1:** Patient Information.

| Patient Sex | Patient ID | Patient Age on Day of Sample Collection | Diagnosis |
| --- | --- | --- | --- |
| <u>Male</u> | SD2 | 75 | Severe aortic stenosis |
|  | SD3 | 71 | Severe aortic stenosis |
|  | SD8 | 74 | Severe aortic stenosis |
|  | SD15 | 82 | Severe aortic stenosis |
|  | SD16 | 79 | Severe aortic stenosis |
|  | SD19 | 84 | Severe aortic stenosis |
|  | SD20 | 82 | Severe aortic stenosis |
|  | SD22 | 78 | Severe aortic stenosis |
|  | SD25 | 64 | Healthy |
|  | SD31 | 78 | Severe aortic stenosis |
|  | SD32 | 77 | Severe aortic stenosis |
|  | SD37 | 87 | Severe aortic stenosis |
|  | SD39 | 65 | Healthy |
|  | SD40 | 76 | Healthy |
| <u>Female</u> | SD5 | 86 | Severe aortic stenosis |
|  | SD10 | 82 | Severe aortic stenosis |
|  | SD13 | 84 | Severe aortic stenosis |
|  | SD26 | 85 | Severe aortic stenosis |
|  | SD27 | 84 | Severe aortic stenosis |
|  | SD30 | 86 | Severe aortic stenosis |
|  | SD34 | 69 | Severe aortic stenosis |
|  | SD35 | 90 | Severe aortic stenosis |
|  | SD36 | 89 | Severe aortic stenosis |
|  | SD41 | 72 | Healthy |
|  | SD42 | 68 | Healthy |
|  | SD44 | 83 | Severe aortic stenosis |
|  | SD45 | 87 | Severe aortic stenosis |
|  | SD46 | 89 | Severe aortic stenosis |

## Notes

### Competing Interest Statement

The authors have declared no competing interest.

